# A polymeric, PDMS-based large-scale skull replacement suitable for optical and mechanical access for long-term neuronal imaging, electrophysiology, and optogenetics

**DOI:** 10.1101/2025.04.15.648966

**Authors:** Kengo Takahashi, Gerjan Huis in ‘t Veld, Davide Benedetti, Jun-Ying Wang, Samuel Pontes Quero, Rafael Yuste, Cyriel M.A. Pennartz, Umberto Olcese

**Affiliations:** Swammerdam Institute for Life Sciences, University of Amsterdam, Amsterdam, NL; Amsterdam Brain and Cognition, University of Amsterdam, Amsterdam, NL; Department of Translational Research and of New Surgical and Medical Technologies, University of Pisa, Pisa, Italy; NeuroTechnology Center, Department of Biological Sciences, Columbia University, New York City, NY, USA

## Abstract

Many techniques to record and manipulate neuronal activity across large portions of the vertebrate brain, such as widefield and two-photon calcium imaging, electrophysiology, and optogenetics, are now available. However, few effective approaches enable both optical and mechanical access to the brain. In this work, we offer an in-depth guide for synthesizing, implanting, and using polydimethylsiloxane (PDMS) windows as skull replacements for chronic optical neuronal imaging. Furthermore, we provide instructions to perform viral injections and multi-site silicon probe implantation.

Graphical abstract

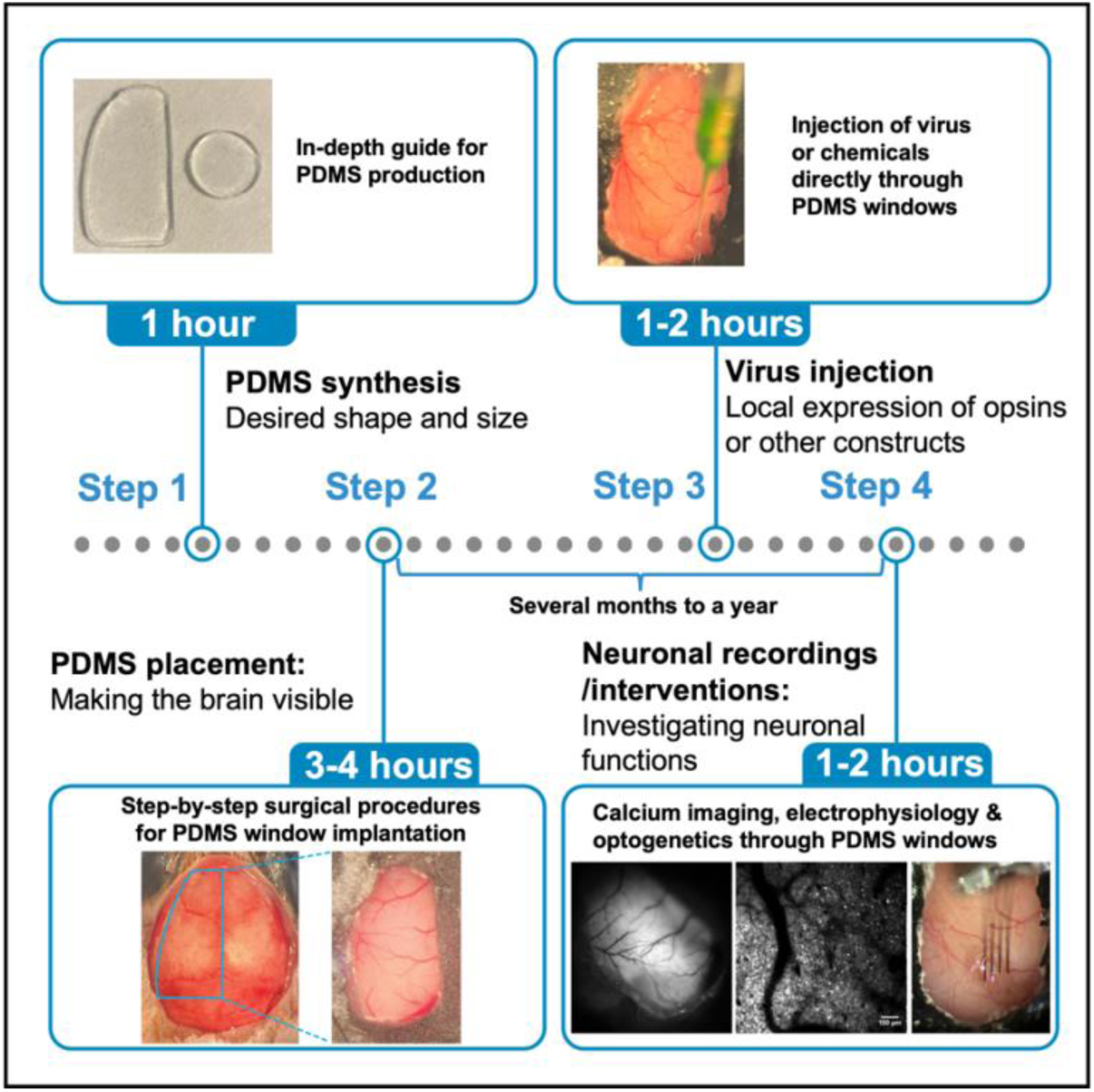

## Before you begin

Recent advancements in neural activity recording techniques, such as calcium imaging and electrophysiology, as well as in neural control methods like optogenetics, have significantly improved our ability to examine the interactions of neuronal dynamics across multiple brain regions during perceptual tasks^1–4^. These multimodal neuronal imaging approaches are particularly relevant in studies aimed at identifying and manipulating specific neuronal populations involved in sensory-motor and cognitive transformations, a process that involves interactions between multiple areas across the whole cortex such as conscious visual processing^5–7^. To perform neural imaging across the whole cortex, the skull needs to be made transparent. Glass windows have long been used to enable functional, neuron-level imaging^8^, but their rigidity makes them unsuitable for replacing the whole dorsal skull in long-term use over several months. Skull-clearing techniques^9^ do not suffer from this limitation, but, similarly to glass windows, do not allow mechanical access to the brain (e.g., for viral injections) without opening a craniotomy, a procedure which reduces transparency and increases the risk of infections. Finally, 3D-printed skull replacements have emerged to enable multi-site silicon probe recordings^1,10^, but are not currently suitable for large-scale imaging. Flexible polymeric windows, e.g. made of polydimethylsiloxane (PDMS), potentially allow to overcome the limitations of the previously mentioned approaches, but standardized protocols to employ them are currently not available. To address this, we provide comprehensive, step-by-step instructions for synthesizing biologically compatible PDMS for use in neuronal imaging. We detail the surgical procedure for PDMS implantation, followed by methods for viral or chemical injections through the PDMS window. Finally, the protocol demonstrates the long-term stability for several months to a year and quality of the implanted PDMS windows via examples of how widefield and two-photon neural imaging can be achieved and integrated with multiple techniques, including electrophysiology and optogenetics. This protocol is primarily designed for mice. However, the presented technique is adaptable to neuronal imaging of other species, such as rats^11^, ferrets^12^, and birds^13^, by adjusting the size and shape of the PDMS window to suit specific experimental requirements. Thus, this protocol offers a comprehensive potential for investigating brain functions in a variety of neuroscience methodologies.

### Institutional permissions

All animal experiments described here complied to international and national legislation and were approved by the Dutch Commission for Animal Experiments and by the Animal Welfare Body of the University of Amsterdam.

### Key resources table

| REAGENT or RESOURCE | SOURCE | IDENTIFIER |
| --- | --- | --- |
| Antibodies |  |  |
| Alexa Fluor 488 Hydrazide | Thermo Fisher Scientific | A10436 |
| Bacterial and virus strains |  |  |
| AAV8-hSyn1-Jaws_KGC_tdTomato_ER2-WPRE-bGHp(A) | Viral Vector Facility - Universität Zürich | v988-8 |
| Chemicals, peptides, and recombinant proteins |  |  |
| Baytril | Bayer Animal Health GmbH | E110564 |
| Carprofen | Dechra | REG NL 112210 |
| Eye ointment (LubriThal) | Dechra | 321/00/12PUVPT |
| Lidocaine 2% | Dechra | REG NL 8632 |
| Saline (0.9% sodium chloride) | Fresenius Kabi | 1310221 |
| Simplex Rapid | Kemdent | ACR920 |
| Betadine | Mylan | RVG 01331 |
| Dexamethasone | Schnelldorf Distribution | PHR1768-1G |
| Dental cement | Sun Medical Co.,Ltd | K058E |
| Isoflurane | Zoetis | B506 |
| Experimental models: Organisms/strains |  |  |
| Mouse: Rasgrf2-dCre (JAX #022864) x Ai148D (JAX #030328) | The Jackson Laboratory | JAX #022864 (Rasgrf2-dCre)<br>JAX #030328 (Ai148D) |
| Recombinant DNA |  |  |
| AAV-Syn-Jaws-KGC-tdTomato-ER2 | Addgene (Chuong et al.<sup>14</sup>) | Plasmid #153535 |
| Software and algorithms |  |  |
| Matlab-based acquisition software and a pre-processing pipeline for widefield imaging | Couto et al.<sup>2</sup> | https://github.com/churchlandlab/WidefieldImager |
| Open Ephys GUI for Neuropixels Probes | Open ephys | https://github.com/open-ephys/plugin-GUI |
| Kilosort3 | Pachitariu et al.<sup>15</sup> | https://github.com/MouseLand/Kilosort/tree/v3.0 |
| suite2p | Pachitariu et al.<sup>16</sup> | https://github.com/MouseLand/suite2p |
| ThorImageLS 4.3 | Thorlabs | https://www.thorlabs.com/newgrouppage9.cfm?objectgroup_id=9072#ad-image-0 |
| Other |  |  |
| 0.3-ml syringes (BD Micro-Fine + Demi) | BD | 324826 |
| 25 gauge needle (BD Microlance 3) | BD | 300600 |
| 30 gauge needle (BD Microlance 3) | BD | 304000 |
| Vacuum pump | BILTEC (Polyestershoppen BV) | BT115VM |
| Blue bandpass filter | Chroma | ET470/40x |
| Gauze | Cutisoft | 45846-00 |
| SYLGARD™ 184 Silicone Elastomer Kit | Dow Inc. | 1060040_S1 |
| Nanoject II™ Microinjectors | Drummond Scientific | # 490007-164 |
| Nanoject Glass Capillaries | Drummond Scientific | 3-000-203-G/X |
| Flexdym 250 µm | Eden Tech | M-FS |
| Green Bandpass Filter | Edmund Optics | 86-963 |
| Green Dichroic Filter (50mm Square 45°) | Edmund Optics | 47-950 |
| Dental drill | Eickemeyer |  |
| Isoflurane vaporizer | Eickemeyer | 213261 |
| Petri dish (90 x 16 mm) | Eurofysica | 016610 |
| Absorbable haemostatic gelatin sponge (Spongostan Dental) | Ferrosan Medical Devices | MS0005 |
| Fine scissors | Fine Science Tools | 14094-11 |
| Forceps | Fine Science Tools | 11251-35 |
| Petri dish (145 x 20 mm) | Greiner Bio-One | 639102 |
| 0.5mm diameter drill tip | Hager & Meisinger GmbH | 310204001001005 |
| 0.9mm diameter drill tip | Hager & Meisinger GmbH | 310204001001009 |
| Cotton swab | Heinz Herenz | TM1301105 |
| Super glue (Pattex ULTRA GEL super glue) | Henkel | 5410091669058 |
| Neuropixels 1.0 probe | IMEC (Jun et al.<sup>17</sup>) | PRB_1_4_0480_1_C |
| Neuropixels 2.0 probe | IMEC (Steinmetz et al.<sup>18</sup>) | NP2014 |
| Biopsy punch | Kai Medical | BP-15F |
| Vacuum Dome (8 Liter) | Kartell (Polyestershoppen BV) | N/A |
| Stereotaxic device (Harvard Apparatus U-frame Stereotaxic Instrument) | Multi Channel Systems | 751810 |
| Nikon AF-D 105MM F/2.0 DC Occasion | Nikon | 018208019328 |
| 5-ml syringes | Nipro | SY3-5LC-GEC |
| 1-ml syringes | Servoprax GmbH | I3040200 |
| Sterile absorbent swabs (Sugi sponge points) | Questalpha GmbH & Co.KG | 31602 |
| Ultra Gel (thick glue) | Pattex | 5410091669058 |
| Twinsil (two-component silicone-glue) | Picodent | 13001000 |
| 85mm F1.4 Aspherical Fixed Lens | Rokinon | 85M-C |
| pco.edge 5.5 USB sCMOS Camera | PCO AG (SDVision) | pco.edge 5.5 M-AIR-CH-BLU-P |
| Femtosecond laser (920nm) | Spark Lasers | ALCOR 920 |
| Collimator with Ø2" Lens, AR Coating: 350 - 700 nm | Thorlabs | SM2F32-A |
| Scalpel Blades | Swann-Morton | 0205 |
| Mounted LED (470 nm) | Thorlabs | M470L5 |
| Multiphoton Imaging Microscope | Thorlabs | Bergamo II |
| Nikon Plan Fluorite Water Dipping Objective (16X) | Thorlabs | N16XLWD-PF |
| T-Cube LED Driver, 1200 mA Max Drive Current | Thorlabs | LEDD1B |

### Materials and equipment setup

This protocol employs a standardized neural recording system, featuring Neuropixels probes for electrophysiological recordings^17,18^, a pco.edge 5.5 USB sCMOS camera (PCO AG) for widefield calcium imaging^2^, and a multiphoton imaging microscope (Bergamo II, Thorlabs) for two-photon imaging.

## Step-by-step method details

### PDMS synthesis protocol

**Timing: [1h (2-3 days waiting)]**

1. Add 40 g of Sylgard 184 polymeric base (Dow Inc.) into a plastic Petri dish (Figure 1a, left). In this example, a 145 × 20 mm Petri dish was used, but the size is not crucial at this stage, as the primary goal is to mix the two components. A larger dish can be used for larger batches, while a smaller one is suitable for smaller portions. To prevent spills and simplify cleanup, it is advisable to place a layer of plastic wrap on the scale before pouring the silicone elastomer.
2. Add 4 g of Sylgard 184 curing agent (Dow Inc.), maintaining a 10:1 ratio between the base and curing agents (Figure 1a, middle).
3. Mix vigorously using a sterile stick for 5-10 minutes until the components are homogeneously blended (Figure 1a, right). During this process, a significant amount of air bubbles will form in the mixture (Figure 1c, left).
4. Apply a vacuum (pressure: 0.05 mbar at room temperature) for 5-10 minutes, then slowly open a valve to reintroduce air into the chamber (Figure 1b, right). The bubbles in the liquid will rise to the surface (Figure 1b, left) and then disappear. **Critical:** Rapid valve opening may disrupt the mixture.
5. Repeat Step 4 multiple times until the majority of air bubbles (∼95%) has been removed.
6. Transfer the mixture to another plastic Petri dish to achieve the desired thickness, which was targeted between 300–400 μm. This range was chosen to ensure that the PDMS was not too thin, which could lead to structural damage and potential harm to the brain, while also maintaining sufficient thinness for effective imaging and ease of making an opening in the PDMS window for electrophysiology later (see the expected outcomes section, Figure 4a and 5a for details). For instance, 2.89 g of the mixture was placed in a new Petri dish with a diameter of 8.5 cm, yielding a theoretical PDMS thickness of approximately 500 μm. However, the final PDMS thickness in the area intended for use is typically about 100 μm thinner than the theoretical value, due to the accumulation of PDMS along the edges of the dish, caused by its high viscosity during solidification. Thus, adjustments may be required to achieve a more precise thickness. **Critical:** The adopted container must be smooth and flat, such as a Petri dish, to ensure the surface of the PDMS also becomes smooth, preventing optical distortions.
7. Repeat Step 4 until all visible air bubbles are eliminated (Figure 1c, right). If air bubbles still remain, extend the vacuum duration beyond 10 minutes. Optionally, leave the PDMS in the vacuum for 30 minutes to 1 hour, and repeat the process as necessary until the bubbles are fully eliminated.
8. Leave the Petri dish on a flat surface and allow the PDMS to solidify fully (∼2 days). Alternatively, the process can be accelerated by applying heat using an oven. **Critical:** It is crucial to select a location with minimal slope, as gradients can result in uneven PDMS thickness across different areas of the petri dish. Once the PDMS has fully solidified, it is important to measure its thickness to determine if it is consistent. If not, use PDMS from regions where the thickness falls within the desired range (ideally between 300-500 μm).
9. On the day of PDMS placement surgery, cut the solidified PDMS into the desired shape – e.g., circular, hemispherical, etc. (Figure 1d). Using a mold, such as a biopsy punch, can assist with shaping, though a surgical knife or scissors can also be used. In the example illustrated in Figure 1d (bottom left), the mold was fabricated by milling after being designed for 3D printing. As described in this procedure, 40 g of Sylgard 184 polymeric base and 4 g of Sylgard 184 curing agent are used to prepare 11 Petri dishes (Dimensions: 90 x 16 mm, with a measured diameter of 85 mm for the plate used to pour PDMS) of PDMS. From each Petri dish, approximately 50 PDMS windows can be obtained.

**Figure 1.**
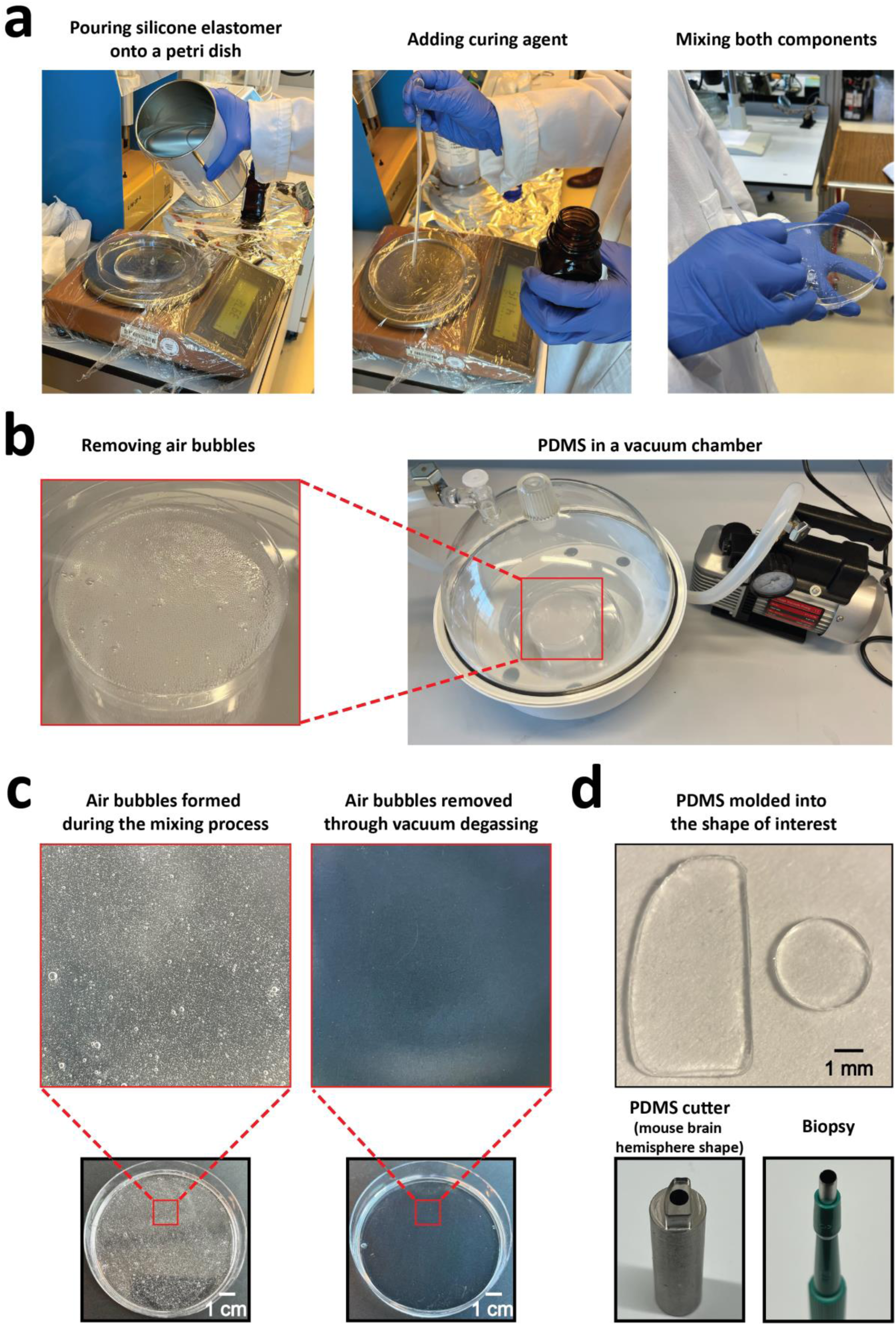
PDMS fabrication process. **(a)** Weighing the silicone elastomer (left) and curing agent (middle), followed by mixing both components (right). **(b)** Removing air bubbles using a vacuum chamber. **(c)** Side-by-side comparison of PDMS before (left) and after (right) bubble removal. d) PDMS molded into specific shapes, such as a hemisphere of the mouse brain using a PDMS cutter (left) and a circle using a biopsy (right).

### Surgical protocol for PDMS placement

**Timing: [∼ 3-4 h]**

1. Anesthetize the animal using isoflurane (3.0–4.0% for induction) and maintain anesthesia at a concentration of 1.5–2.0% throughout the surgical procedure.
2. Trim the fur at the surgical site with a hair clipper (Figure 2a.1) and disinfect the area using 70% ethanol followed by Povidone-iodine solution (Figure 2a.2 and 3).
3. Apply ophthalmic ointment (Figure 2a.4) to the eyes to prevent dryness and administer lidocaine to the shaved skin for local anesthesia (Figure 2a.2 and 5).
4. Administer carprofen (5–10 mg/kg) via subcutaneous injection for analgesia (Figure 2a.6).
5. Administer dexamethasone (8 mg/kg) via intramuscular injection into the hind limb to reduce brain inflammation prior to performing the craniotomy (Figure 2a.7 and 2c.7).
6. Expose the skull by incising the skin with surgical scissors (Figure 2a.8) and secure the skin laterally to the skull using Vetbond (Figure 2a.9 and 2b.1). A surgical scalpel (Figure 2a.10) may be used to assist in retracting the skin; however, avoid cutting the skin or muscle with the scalpel.
7. Place a PDMS window on the skull and use a black marker to draw a line indicating the area for drilling (Figure 2b.2). Ensure the PDMS window is smaller than the craniotomy.
8. Attach a headbar to the skull using super glue (Figure 2a.11, 12, and 2b.3). The headbar should be positioned slightly towards the left side of the skull if securing the entire left hemisphere of the cortex is required. To achieve this, apply glue only to the upper right side of the headbar. The lower left part of the headbar should not contact the skull and should remain elevated. Ensure the headbar is parallel to the head’s orientation, as a tilted headbar may alter the animal’s visual field during visual detection tasks later. In the example shown in Fig. 2b.2, the approximate center of the visual cortex (A/P = 4.0 mm, M/L = 2.5 mm) is marked with a black dot.
9. Apply dental cement to the headbar (Figure 2a.13 and 2b.4).
10. At this step, roughly one hour into the surgery, administer a subcutaneous injection of saline (Figure 2b.14) to prevent hypovolemia. Continue administering saline every hour for the remainder of the surgery.
11. Drill along the previously drawn line until the skull is thin enough to move when touched (Figure 2b.5). Start with a 0.9 mm drill bit (Figure 2a.15) to outline the area, then switch to a 0.5 mm bit (Figure 2a.16) to refine and thin any remaining thick areas. Be sure to smooth the outer edge of the drilled region to avoid leaving any skull fragments attached post-removal.
12. Ensure that the metal plate intended to press the PDMS window later can be properly positioned and angled within the craniotomy (Figure 2b.6).
13. First apply cold saline, then carefully flip the drilled area of skull using tweezers (Figure 2a.17) to expose the brain (Figure 2b.7). Once the skull has been flipped, wipe the edge of the craniotomy with a triangular sponge (Figure 2a.18). If there is bleeding, apply cold saline again. It is critical to have cold saline, cotton swab, and absorbable haemostatic gelatin sponges (Figure 2a.19 and 20) prepared in advance before flipping the skull in case of bleeding.
14. If bleeding occurs, wait for it to stop before proceeding. Then, place the PDMS window onto the brain (Figure 2b.8), ensuring it is clean. If not, wash the PDMS window with isopropyl alcohol and dry it before placement.
15. Press the PDMS window until it is either at the same level as the skull or slightly below it (Figure 2b.9). **Critical**: Do not apply excessive pressure to the PDMS window, as this can slow the animal’s breathing rate, which can be fatal. Continuously monitor the animal’s respiration while pressing the PDMS window.
16. Fill the gap between the PDMS window and the skull with thick glue (Figure 2a.21), ensuring that the glue does not adhere to the metal plate used to press the PDMS window (Figure 2b.10).
17. Slowly remove the metal plate that was used to hold the PDMS window in position (Figure 2b.11).
18. Apply dental cement over the glue, around the edge of the PDMS window, and on the headbar (Figure 2b.12). In this example, black ink was mixed with the dental cement (Figure 2a.21, 22, and 23) for better protection from stray light for calcium imaging.
19. Place a metal ring on top of the headbar and secure it using dental cement (Figure 2c.1 and 2). Once the cement has fully hardened, place a black cover over the ring (Figure 2c.1 and 3). This procedure protects the PDMS window by minimizing its exposure during housing. If a metal ring and black cover are unavailable, an alternative approach is to cover the PDMS window (Figure 2c.4) with dried gauze (Figure 2c.5) and secure it with a metal plate (Figure 2c.6). Ensure that the PDMS window is dry before placing the gauze.

**Figure 2.**
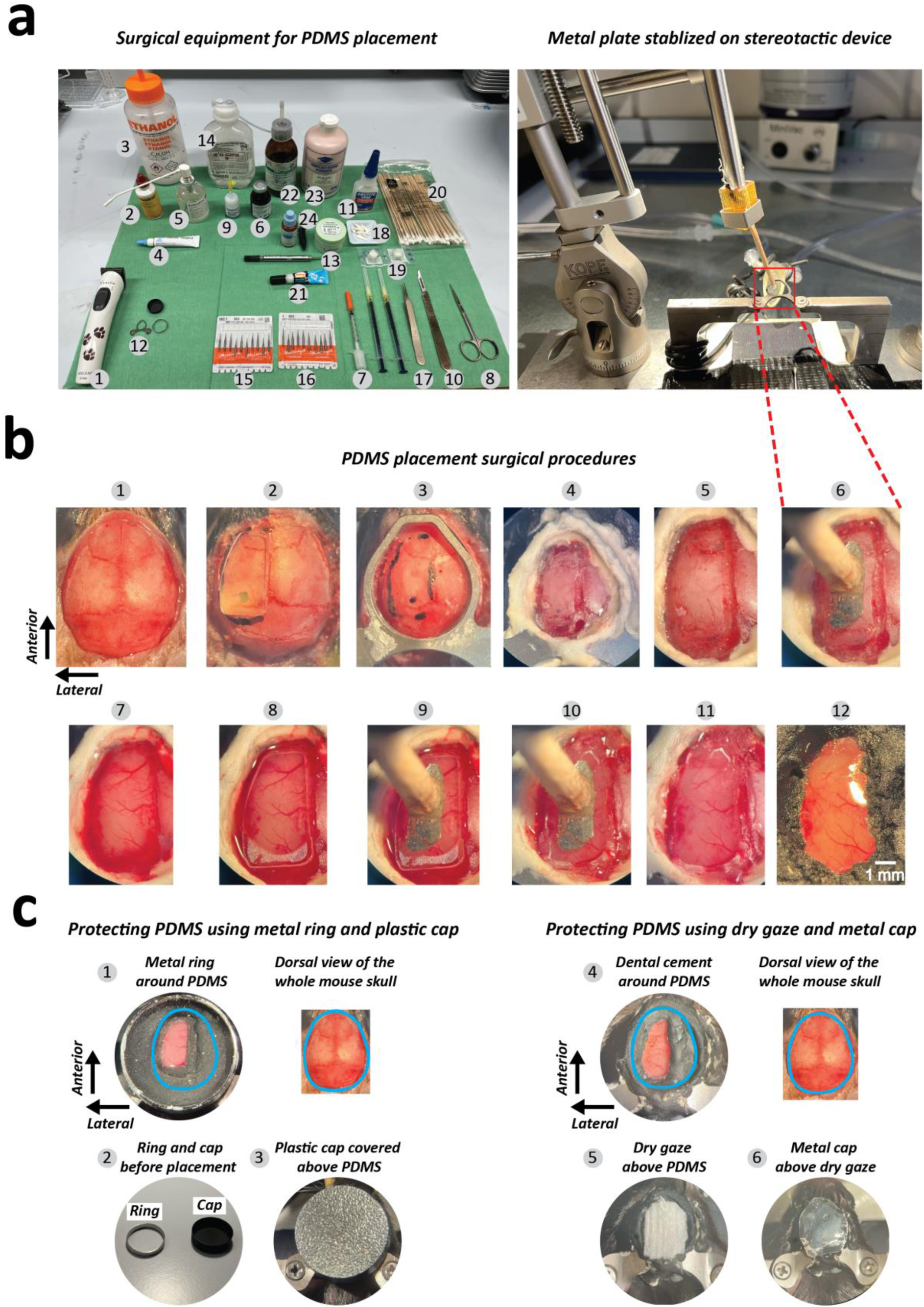
Surgical procedure for PDMS implementation. **(a)** Surgical Setup and Equipment: Displays the surgical instruments (left) and the setup (right) required for PDMS placement. (1) Hair shaver. (2) Povidone-iodine. (3) 70% Ethanol. (4) Eye ointment. (5) Lidocaine. (6) Carprofen. (7) 0.3 ml syringes. (8) Surgical scissors. (9) Vetbond. (10) Surgical scalpel. (11) Super glue. (12) Head bar. (13) Dental cement. (14) Saline. (15) Drill tip (0.9 mm). (16) Drill tip (0.5 mm). (17) Tweezers. (18) Triangular sponges. (19) Gelatin sponge. (20) Cotton swab. (21) Thick glue. (22) Catalyzer for dental cement. (23) Dental cement. (24). Black ink. **(b)** Surgical Procedure: 1) The skull is exposed by making an incision in the skin. (2) The PDMS window is positioned on the skull, and a reference outline is drawn for drilling. (3) The headbar is temporarily secured using glue. (4) Dental cement is applied over the headbar for stabilization. (5) The skull is drilled and thinned along the marked outline. (6) A metal plate is placed to ensure proper alignment for later PDMS compression. (7) The skull is removed to expose the brain. (8) The PDMS window is positioned over the exposed brain. (9) The PDMS window is pressed down until it aligns with the level of the skull. (10) The gap between the PDMS window and dental cement is sealed with thick glue. (11) The metal plate holding the PDMS window is removed. (12) A final layer of dental cement, mixed with black ink, is applied over the headbar, glue, and a small portion (∼0.1 mm) of the PDMS window. **(c)** PDMS window protection: (1) Uncovered view of the PDMS with metallic ring, and dorsal view of the whole mouse skull. The ring is secured with dental cement. The original position of the mouse skull is indicated in blue. (2) A protective ring and cap are used to safeguard the PDMS. (3) The PDMS is covered with the cap. (4) Uncovered view of the PDMS and brain without the ring and cap, and dorsal view of the whole mouse skull. (5) A layer of dry gauze is placed over the PDMS for protection. (6) A metal plate is secured over the gauze for additional protection.

### Virus or chemical injection protocol through the PDMS window

**Timing: [∼ 1-2 h]**

1. Use a puller to fabricate a thin tip (∼100 μm diameter) on a glass capillary for virus or chemical injection.
2. Fill the pulled glass capillary with the solution containing the necessary virus or chemicals, and attach it to the microinjector (Figure 3a). In this example, Adeno-Associated Viruses (AAV) were used, and Alexa Fluor 488 dye was included to visualize and verify the success of the injection.
3. Make a small incision on the surface of the PDMS window at the top of the region of interest (ROI) using a 30G needle, where the virus will be injected (Figure 3b). This incision is necessary to facilitate the entry of the pulled glass capillary into the PDMS window, as the high surface tension of PDMS can impede insertion. **Critical**: The incision does not need to extend all the way to the bottom of the PDMS window; a small superficial incision is sufficient.
4. Insert the pulled glass capillary into the small incision (Figure 3c).
5. Lower the capillary until it makes contact with the surface of the dura mater. When the capillary touches the dura, the movement of the dura (or brain) becomes readily observable, allowing for the subsequent calculation of injection depth. This point is designated as 0 mm on the dorsal-ventral axis. If the brain is not clearly visible due to insufficient transparency of the PDMS window, the addition of demineralized water can enhance visualization. **Note:** The use of saline, phosphate-buffered saline (PBS), or tap water may lead to salt deposits on the PDMS window, possibly resulting in a foggy window. Once the PDMS window is contaminated, cleaning can be challenging.
6. Continue to lower the capillary until it punctures the dura mater. Upon puncturing the dura, a rebound effect can be observed as both the dura and brain return to their original positions due to the release of tension.
7. Position a glass capillary at the desired depth for injection.
8. Inject the viral suspension into the ROI and wait for approximately 10 minutes. If additional injections are necessary, repeat the procedure at various sites.
9. Remove the capillary slowly to minimize the risk of backflow of the injected virus.
10. Cover the PDMS window with a plastic cap or dried gauze and metal plate (Figure 2c.3 and 6).

**Figure 3.**
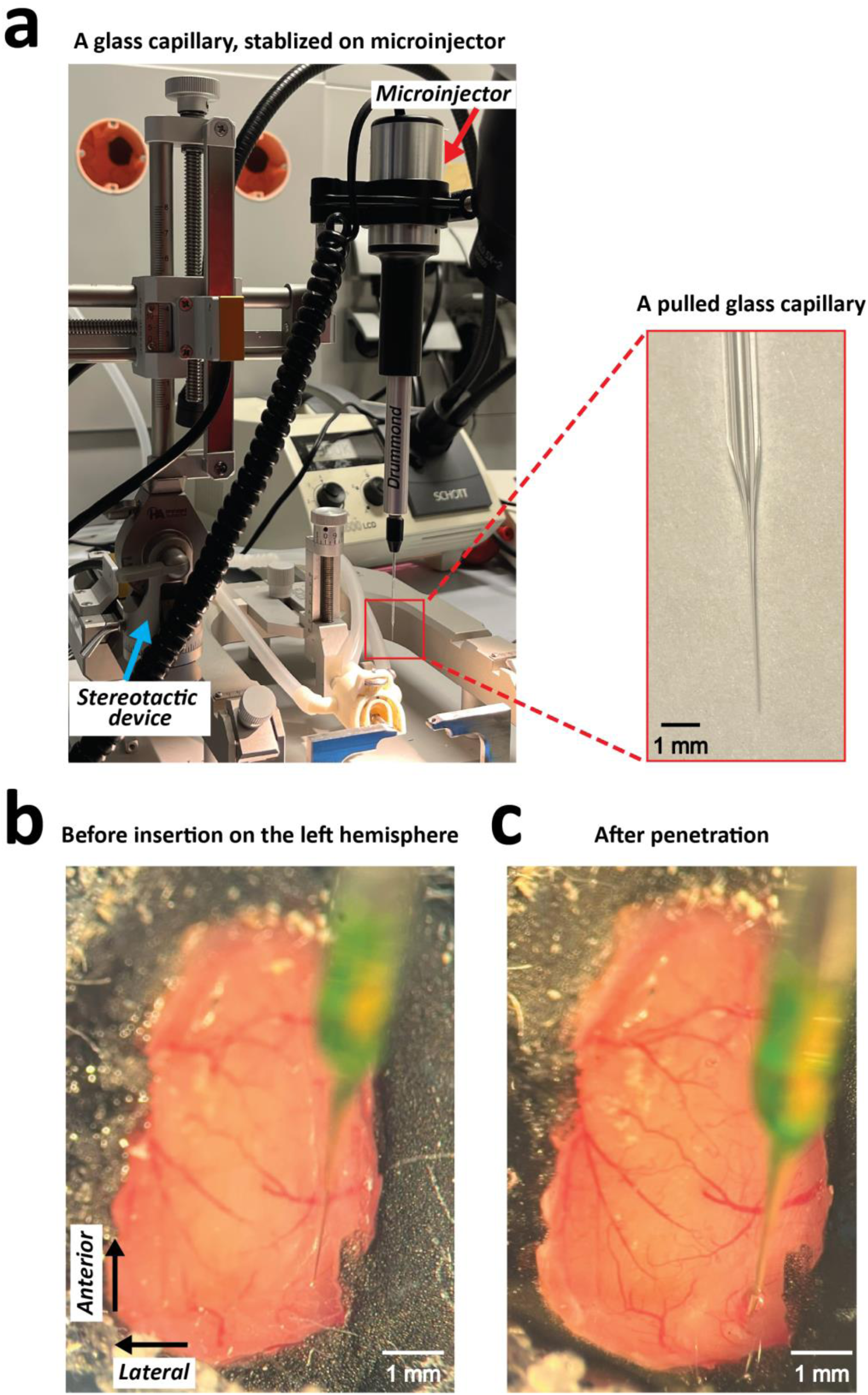
Virus injection procedure on PDMS. **(a)** A glass capillary mounted on a microinjector within a stereotactic device. The enlarged image displays the tip of a glass capillary, with an approximate tip diameter of 100 μm and a base diameter of around 1 mm. **(b)** The capillary positioned above the PDMS prior to virus injection on the left hemisphere of the mouse brain. **(c)** The capillary penetrating the PDMS during the injection process.

### Expected outcomes

This protocol facilitates the detection of neural activity across various cortical regions of interest, leveraging the flexibility of PDMS fabrication in different sizes and shapes. A key advantage of this method is its ability to first monitor neural activity, such as through calcium imaging, in the context of specific experimental paradigms. Then, based on the imaging results, targeted virus or chemical injections can be performed in specific brain regions for precise manipulation, as the PDMS window allows easy penetration by a glass capillary, enabling precise viral or chemical injections. Notably, the PDMS window remains optically clear for several months (and up to a year) using this protocol. If deterioration occurs, a replacement procedure can be undertaken. To illustrate the capabilities of this method, we present the results of two experiments demonstrating reliable neural activity detection using widefield and two-photon calcium imaging, as well as the feasibility of electrophysiological recordings combined with optogenetics through the PDMS window.

### Widefield or two-photon calcium imaging recording through PDMS window

By utilizing a transgenic animal expressing a calcium indicator, excitatory neuronal activity can be monitored through a PDMS window using both widefield and two-photon calcium imaging. For instance, widefield imaging was used to capture a clear view of the brain and its vasculature of a mouse over 10 months after implantation (Figure 4a). Neural activity was detected in the visual cortex of GCaMP6f-expressing mice (Rasgrf2-dCre x Ai148D) in response to visual stimuli, such as a moving bar (Video 1) or a Gabor patch (Figure 4b). While this example demonstrates the feasibility of detecting neural responses to visual stimuli, the experimental design can be adapted to study different sensory modalities, such as auditory stimuli, for which activation of the auditory cortex and distinct neuronal activity patterns would be expected. Since the PDMS window covers one entire cortical hemisphere, this enables simultaneous investigation of multiple cortical regions. Furthermore, neuronal activity can be also studied through the PDMS window using two-photon microscopy (Figure 4d and Video 2), for which we show recordings obtained one month (Figure 4c) and seven months (Figure 6b, right) after implantation. This methodology enables researchers to first identify regions of interest using widefield imaging and subsequently perform high-resolution, single-neuron investigations within the same animal using two-photon microscopy, enabling a comprehensive analysis of cortical dynamics.

**Figure 4.**
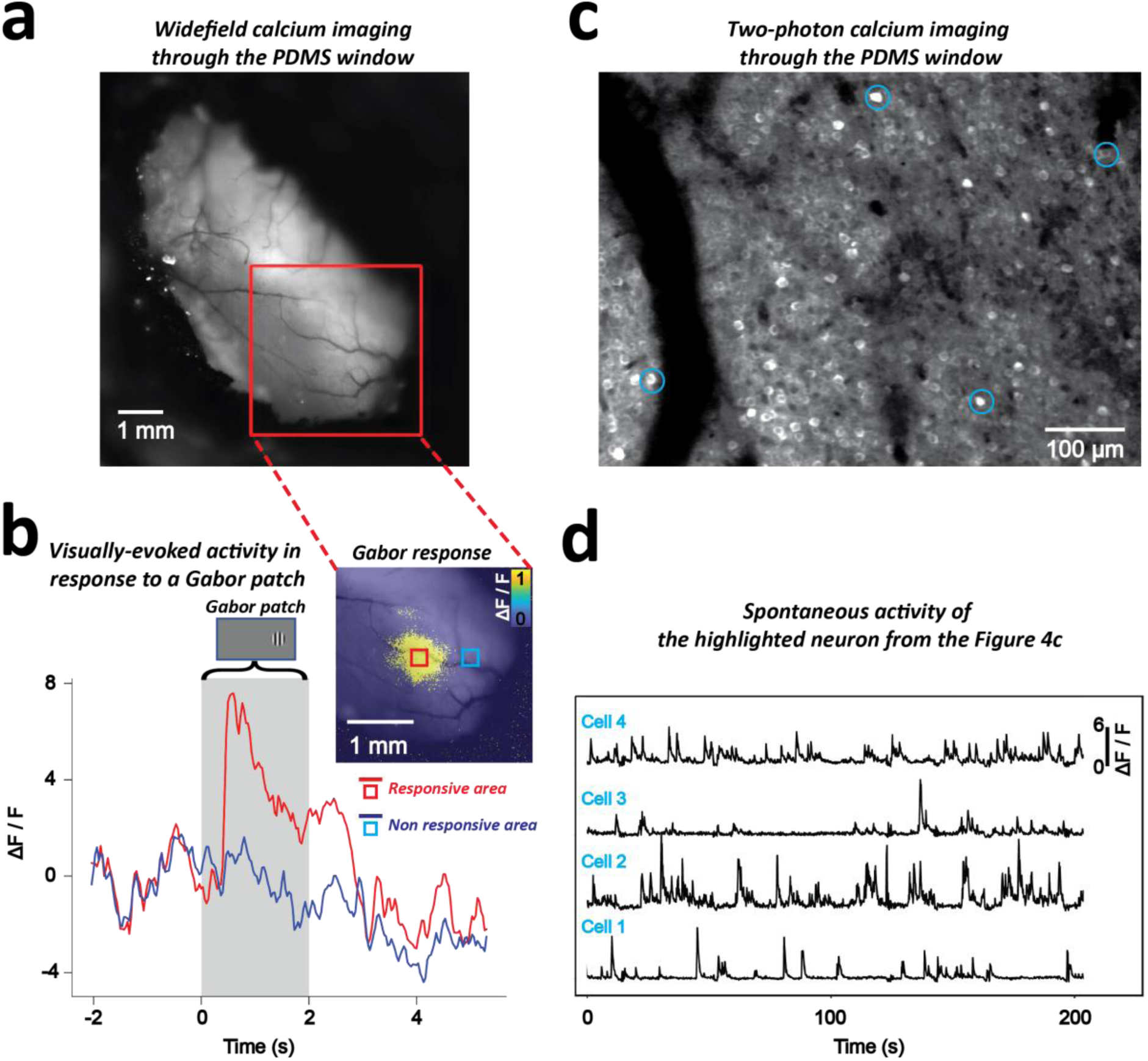
Widefield and two-photon calcium imaging through a PDMS window. **(a)** Widefield calcium imaging through the PDMS window at the 43rd week after the PDMS implantation using a widefield imaging camera, shown with the region of interest (red square) for calcium imaging in the left visual cortex. **(b)** Visually-evoked activity in response to the presentation of a Gabor patch. Left, top: An example of Gabor patch shown on a gray background in the right (contralateral) visual field. Right, top: Imaged neuronal response (ΔF/F) to the Gabor patch in the visual cortex, averaged over 40 trials. Each trial consisted of 2s pre-stimulation baseline, 2s visual stimulation, and 3s post-stimulation. Bottom: ΔF/F trace over time, averaged across 40 trials, from the neuronal region of interest indicated by the blue and red square in the top-right image. The red square marks the region that responded to the Gabor patch presentation, with its averaged neural response shown by the red line. The blue square marks the non-responsive region, with the blue line representing its corresponding neural activity. **(c)** Two-photon calcium imaging over the PDMS window. Example neurons selected for further analysis are marked with a blue circle. **(d)** Spontaneous activity of the highlighted neuron from the Figure 4c. The ΔF/F traces represent the normalized changes in fluorescence over a period of 200 seconds for the neuron.

**Figure 5.**
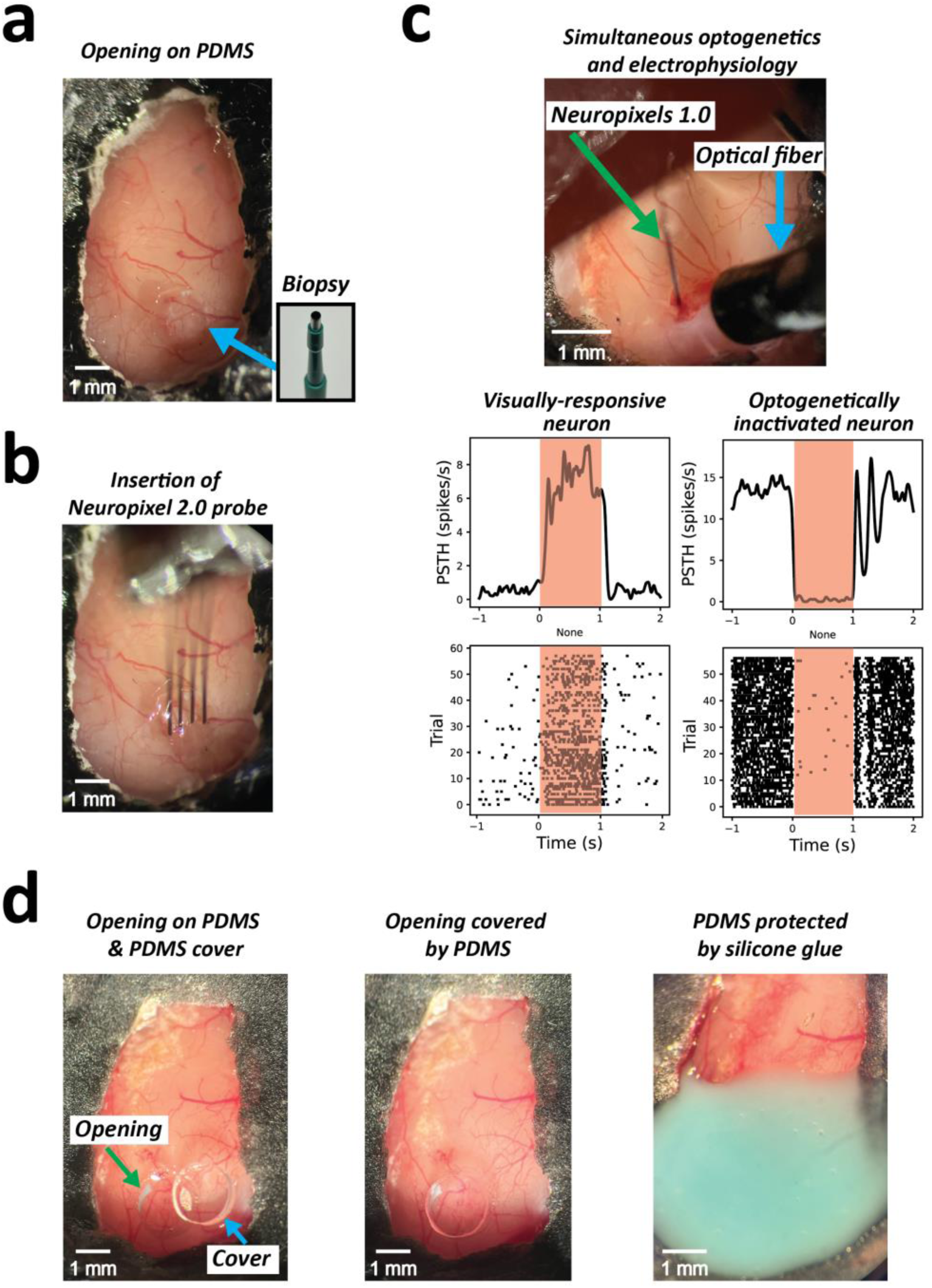
Optogenetic manipulation and electrophysiological recording via the PDMS window **(a)** Example of an opening performed on a PDMS window using a biopsy punch. **(b)** Insertion of a Neuropixels 2.0 probe through the window opening. **(c)** Simultaneous optogenetics and electrophysiology over the PDMS window. A Neuropixels 1.0 probe was used for electrophysiological recording over the PDMS window. The bottom left panel illustrates neural responses to visual stimuli, while the bottom right panel depicts neuronal inhibition. Red shaded areas indicate the period of sensory or optogenetic stimulation, respectively. **(d)** Sealing procedure following electrophysiological recordings. The left panel displays an exposed opening in the PDMS with a corresponding PDMS cover positioned adjacent to it. The middle panel displays the opening after being sealed with the PDMS cover. The right panel displays the application of two-component silicone glue (Twinsil) over the sealed area to ensure additional protection.

**Figure 6.**
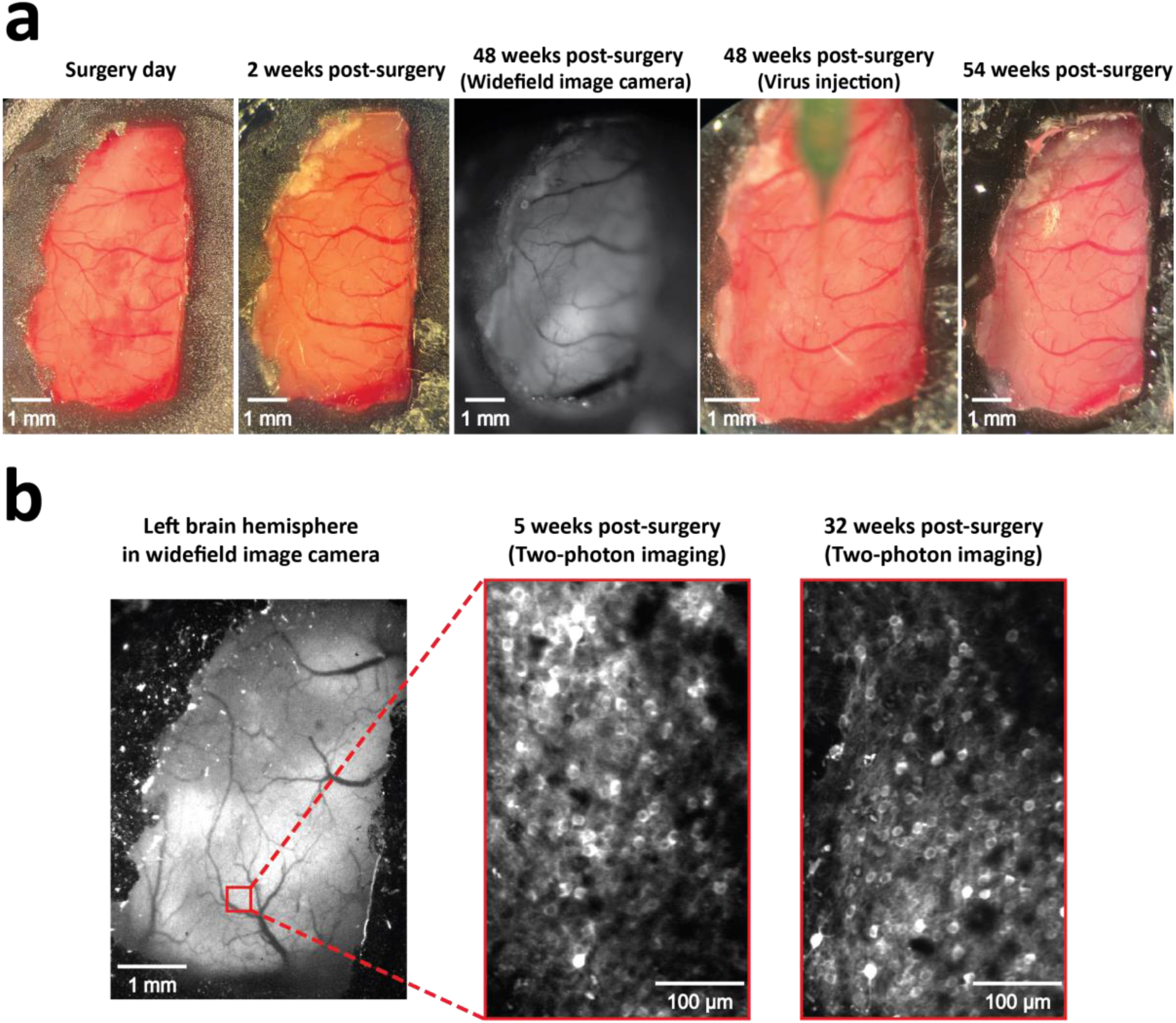
(a) Longitudinal monitoring of the PDMS window in the same animal from the day of surgery up to one year. The images, from left to right, depict: the day of surgery, 2 weeks post-surgery, 48 weeks post-surgery on the day of calcium imaging using a widefield camera, 48 weeks post-surgery during virus injection, and 54 weeks post-surgery. **(b)** Longitudinal assessment of two-photon calcium imaging. Left: Image of the entire left hemisphere of the cortex by widefield imaging camera, with the red square highlighting the region of interest for two-photon imaging. Middle: Two-photon calcium imaging at 5 weeks post-surgery. Right: Two-photon calcium imaging at 32 weeks post-surgery.

**Note**: In the example illustrated in Figure 4a, a Gabor patch was presented on a gray screen with the following parameters: 40 trials, a diameter of 12 degrees, static gratings at a spatial frequency of 0.04 cycles per degree, 100% contrast. The images were captured using a widefield imaging camera with a sampling frequency of 30 Hz and twofold pixel binning. Blue (470 nm, 5 mW, 15 Hz) and violet (405 nm, 5 mW, 15 Hz) LED light were alternately applied to the sample at each frame. In Video 1, a moving bar was used as the visual stimulus, with the following parameters: 25 trials, a stimulus speed of 10 cm/s, a flicker frequency of 6 Hz, a baseline duration of 3 seconds, and a target frame rate of 60 Hz. The images were also captured using a widefield imaging camera with a sampling frequency of 30 Hz and twofold pixel binning, while blue (470 nm, 5 mW, 30 Hz) LED light was applied. For both examples, a screen with a width of 36 cm and a height of 21 cm was situated 11 cm away from the eye to the nearest point of the screen in the right visual field. In the example illustrated in Figure 4c and d, two-photon calcium imaging was conducted using a large field 2P microscope equipped with a 16× 0.8 NA objective (Nikon) and a digital zoom of 2×. Imaging was performed with an 80 MHz ultrafast pulsed laser at a fixed wavelength of 920 nm (Spark Lasers). Frames were acquired at a rate of 7.7 Hz using ThorImageLS software at a resolution of 1,024 × 1,024 pixels, covering a field of view of 973.54 μm × 973.54 μm. Preprocessing of two-photon calcium imaging was performed using the automated segmentation algorithm in Suite2p for neuron detection.

### Optogenetics and electrophysiology recording through PDMS window

Following calcium imaging to identify the region of interest (ROI) for investigation through the PDMS window, two additional methodologies can be further employed. The first approach involves electrophysiological recordings to examine neuronal spiking activity in the identified ROI. Electrophysiology serves as an alternative and as a complementary method to two-photon imaging for assessing neuronal spiking activity, offering substantially higher temporal resolution. A key advantage of PDMS over glass windows is their versatility: researchers can either perform an opening in the PDMS or directly penetrate it with an electrode. However, certain delicate electrodes, such as Neuropixels^17,18^, cannot be directly inserted through PDMS. In such cases, an opening is necessary (Figure 5a). In our study, we successfully inserted Neuropixels 1.0^17^ and 2.0^18^ probes by creating a 1.5 mm opening in the PDMS window using a biopsy punch (Figure 5b,d left). The second approach utilizes the optical transparency of PDMS for optogenetic manipulation. Once calcium imaging has identified the ROI based on the experimental design, viral vectors can be injected into the target area following the protocol detailed in Figure 3. After viral expression, optogenetic stimulation can be effectively applied, as PDMS is transparent, allowing light to penetrate effectively. Moreover, electrophysiology and optogenetics can be combined to causally manipulate and at the same time monitor neuronal responses (Figure 5c, top). As an example, we demonstrate neuronal activation in the visual cortex in response to visual whole-screen gratings (Figure 5c, bottom left) and concurrent neuronal inhibition through optogenetic manipulation (Red light laser with 637 nm wavelength and 8 mW intensity) in a mouse expressing Jaws (mediated by injection of AAV8-Syn-Jaws) (Figure 5c, bottom right). Once the electrophysiology recording is completed, the opening should be sealed by placing a circular piece of PDMS over it (Figure 5d middle). This ensures the maintenance of brain health by preserving moisture and preventing desiccation. The circular piece of PDMS extracted during the biopsy procedure should be reused here, to ensure a tight fit. Otherwise, a new piece can be easily prepared by extracting it from a PDMS plate using a biopsy punch. While a PDMS cover alone may not stay in place, applying a two-component silicone glue (e.g., Twinsil) around the perimeter of the opening can help secure it (Figure 5d right). However, covering the entire PDMS window with silicone should be avoided, as this can make removal difficult for subsequent electrophysiological recordings, due to the adhesive’s strong bonding, potentially causing the entire PDMS window to detach. Thus, it is advised to use a minimal amount of silicone glue.

**Note**: In the example illustrated in Figure 5c, a full-screen grating was presented for 1 second with the following parameters: a spatial frequency of 0.04 cycles per degree, a temporal frequency of 2 Hz, and an inter-trial interval of 3 to 8 second. Consistent with the calcium imaging setup, a screen with a width of 36 cm and a height of 21 cm was also situated 11 cm away from the eye to the nearest point of the screen in the right visual field.

### Performance of PDMS windows over time

While a prior study mentioned that skull regrowth could take place between 5 and 8 months following the implantation of a PDMS window^11^, our protocol enables PDMS windows to retain their optical transparency for extended periods, ranging from several months to a year (Figure 6a). This enabled us to successfully perform both widefield and two-photon calcium imaging over the extended timeframe as indicated by earlier results (Figure 4a and 6b). This extended functionality may be due to the flexible design of our custom-made window and refined surgical techniques (Figure 1 and 2). In our protocol, the PDMS window can be easily shaped into any form or size by using scissors, a biopsy punch, or other comparable instruments (Figure 1). PDMS is flexible and bendable, which is advantageous and well-suited for covering large brain areas with uneven surfaces. When the cranial window is large, rigid materials such as glass may leave gaps between the window and the brain surface due to the brain’s curvature. This curvature can result in insufficient and uneven pressure on the dura during window placement, which can promote dura or skull regrowth over time^19^. In contrast, flexible materials such as PDMS adapt to the brain’s contours, ensuring a snug and secure fit. However, achieving this requires precise surgical techniques rather than simply placing the material on the brain. Otherwise, skull regrowth may occur earlier. Although degradation in the window’s quality is occasionally observed, replacement of the window is feasible with our protocol (see troubleshooting section and Figure 8).

### Comparison with other window types

In calcium imaging studies, researchers typically employ either clear skull procedures or glass windows, depending on the imaging resolution required. Clear skull procedures are commonly used for widefield calcium imaging^1,9,20^, as this technique does not demand high-resolution visualization. In contrast, the visualization of individual neurons requires a highly transparent window, making glass windows the preferred option for chronic two-photon imaging^8,13,21,22^. To provide an assessment of the results that can be expected through the use of PDMS windows, we compared four types of approaches and materials: clear skull, PDMS, Flexdym, and glass (Figure 7 and Table 1). Flexdym (250μm thickness, 15 kN/m tear strength, Eden Tech) is a commercially available material proposed as an alternative to PDMS. This qualitative comparison is based on the experience gained by the authors during the past decade when performing experiment with the different types of windows discussed here. Each window type presented distinct advantages and limitations. In terms of optical clarity, PDMS, Flexdym and glass windows provide the best visibility, allowing clear visualization of both large and small blood vessels. While the clear skull procedure also enables observation of large and some small vessels, its clarity is inferior compared to PDMS, Flexdym and glass windows. Furthermore, the transparency of clear skulls declines over time, often becoming foggy within several months. In contrast, PDMS, Flexdym and glass windows maintain their transparency for longer durations, ranging from several months to a year (Figure 6). The surgical implantation of PDMS, Flexdym and glass windows, however, is more technically challenging and carries greater risk compared to clear skull procedures, as it requires removal of the skull. In contrast, clear skull procedures are entirely non-invasive, relying solely on the use of various glues.

**Figure 7.**
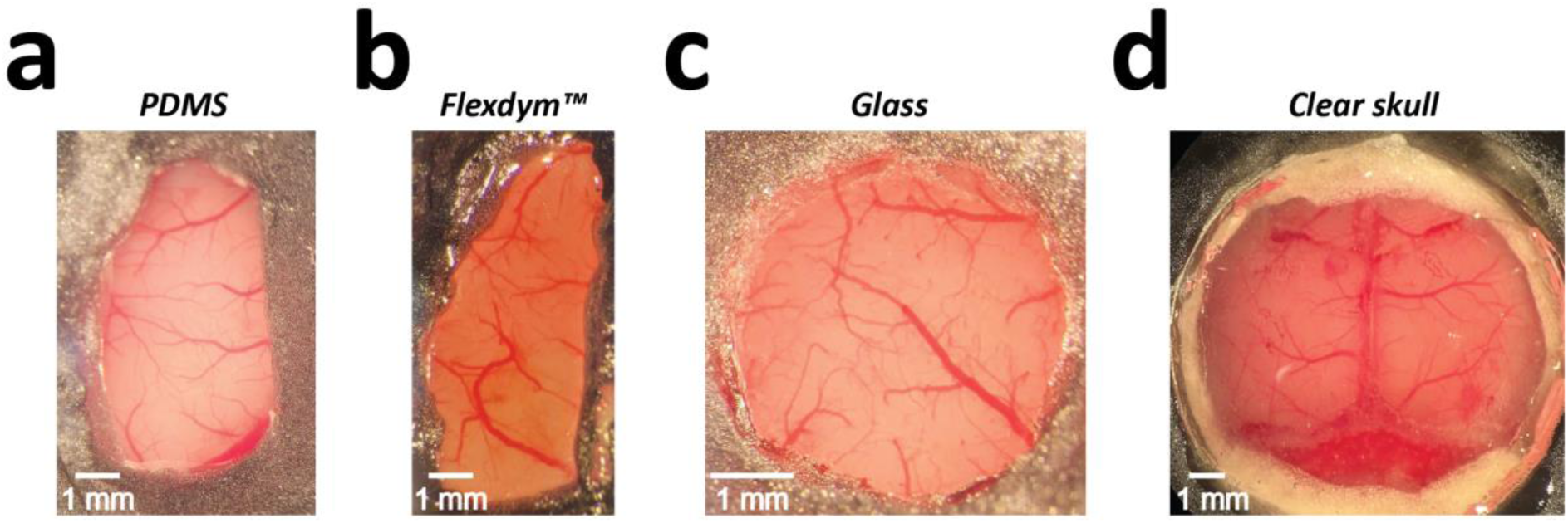
Examples of different window materials: PDMS, Flexdym, glass, and clear skull.

**Table 1.** Qualitative comparison of the optical properties of different window materials: PDMS, Flexdym, glass, and clear skull.

| **Category** | **PDMS** | **Flexdym** | **Glass** | **Clear Skull** |
| --- | --- | --- | --- | --- |
| Clearness | Both large and small blood vessels are clearly visible (Figure 7a) | Both large and small blood vessels are clearly visible (Figure 7b) | Both large and small blood vessels are clearly visible (Figure 7c) | Both large and smaller vessels are visible after surgery but not as transparent as other materials (Figure 7d) |
| Stability Over Time | Relatively stable over a prolonged period | Relatively stable over a prolonged period | Relatively stable over a prolonged period | Quality degrades after several months |
| Surgical Difficulty | Difficult | Difficult | Difficult | Easy |
| Replacement After Degradation | Suitable | Suitable | Suitable | Not feasible |
| Flexibility of Window Shape | Any shape | Any shape, though more challenging to cut | Restricted to predefined shapes (e.g., circular) | n.a. |
| Compatibility with Viral Injection | Compatible | Not compatible | Not compatible | Requires craniotomy |
| Electrophysiology | Requires craniotomy | Not compatible | Not compatible | Requires craniotomy |
| Widefield Calcium Imaging / Optogenetics | Suitable | Suitable | Suitable | Suitable |
| Two-Photon Imaging | Suitable | Suitable | Suitable | Not suitable |

Among PDMS, Flexdym and glass windows, PDMS and Flexdym provide greater versatility in terms of size and shape customization. Glass windows require advance preparation, with predetermined shapes such as circular or rectangular designs. Conversely, PDMS and Flexdym windows can be customized on the day of surgery by cutting the material with scissors, a biopsy punch, or similar tools (Figure 1d). This adaptability makes PDMS and Flexdym particularly advantageous for studies requiring coverage of large brain areas, such as those focused on multisensory integration between the auditory and visual cortices^23^ or sensory-motor transformations^6,7^. In particular, PDMS is particularly well-suited and more advantageous for multimodal approaches that require additional invasive procedures such as viral injections for optogenetics or electrode insertions for electrophysiology. In such cases, using a glass window would require drilling, which is impractical as it disrupts high-quality optical access. Similarly, Flexdym is difficult to penetrate once implanted, since the material is more rigid than PDMS. In contrast, PDMS allows easy insertion of tools such as metal electrodes^11^ or pulled glass capillaries for viral injections (Figure 3c). A notable limitation of PDMS (and of all other window types as well) is its incompatibility with insertion of Neuropixels probe, which are increasingly popular for recording neuronal activity across numerous channels. Neuropixels probes are unable to penetrate PDMS, as they tend to bend and risk breaking. A previously published method involves creating a 3D-printed mold with predefined holes for probe insertion^1^. However, this approach lacks both sufficient optical clarity for calcium imaging as well as flexibility in probe insertion sites, as the holes must be created at predefined insertion locations before window implantation. Thus, this approach restricts the flexibility of insertion sites when the goal is to first identify the insertion areas through calcium imaging and then insert the probe. However, small openings can be made in PDMS using a biopsy punch, enabling Neuropixels probe insertion after identifying the region of interest through calcium imaging (Figure 5a, b).

### Limitations

Neuropixels probes (and other silicon probes) cannot be directly inserted into polydimethylsiloxane (PDMS), whereas more rigid materials such as metal electrodes and glass pipettes can be inserted without issue. This presents a challenge, especially given the remarkable progress made in neurotechnology over the past few years. However, small openings can be made in PDMS windows to enable this type of recordings (see optogenetics and electrophysiology recording through PDMS window section and Figure 5a).

## Troubleshooting

### Problem 1

PDMS not being fully transparent (related to Step 1)

### Potential solution

If inappropriate materials, such as 3D-printed molds, are used for PDMS synthesis, issues may arise. Specifically, two primary problems can occur when PDMS is poured into such molds or other non-smooth materials: (1) incomplete curing, resulting in a partially liquid state, and (2) a lack of full transparency due to the mold’s uneven or rough surface. In our initial attempts, PDMS failed to solidify properly when an unsuitable mold was used. To ensure successful curing and transparency, it is essential to select an appropriate mold material. A petri dish, as recommended in the materials section, provides a smooth, flat surface that ensures PDMS cures properly and remains transparent. Thus, we do not recommend using containers other than Petri dishes, as the surface of alternative containers may not be sufficiently smooth for PDMS synthesis.

### Problem 2

PDMS becomes dirty after being placed over the brain on the day of the surgery (related to Step 2)

### Potential solution

As shown in Figure 8, PDMS can sometimes appear not fully transparent, especially around the area where the metal plate was applied (Figures 2b.9, 2b.10, and 7a). Three potential reasons why PDMS may become dirty are: 1) the metal plate was dirty, 2) PDMS was over-washed or left in ethanol or isopropyl alcohol for an extended period, or 3) glue from the metal plate transferred onto the PDMS during the surgical procedure. To prevent these issues, it is recommended to thoroughly clean the metal plate with isopropyl alcohol before starting the PDMS placement surgery. The PDMS window should also be rinsed with isopropyl alcohol and fully dried before being placed on the brain. However, prolonged exposure to isopropyl alcohol should be avoided as it may negatively affect PDMS transparency. Lastly, avoid allowing glue to come into contact with the metal plate, as it is difficult to clean once it has adhered.

**Figure 8.**
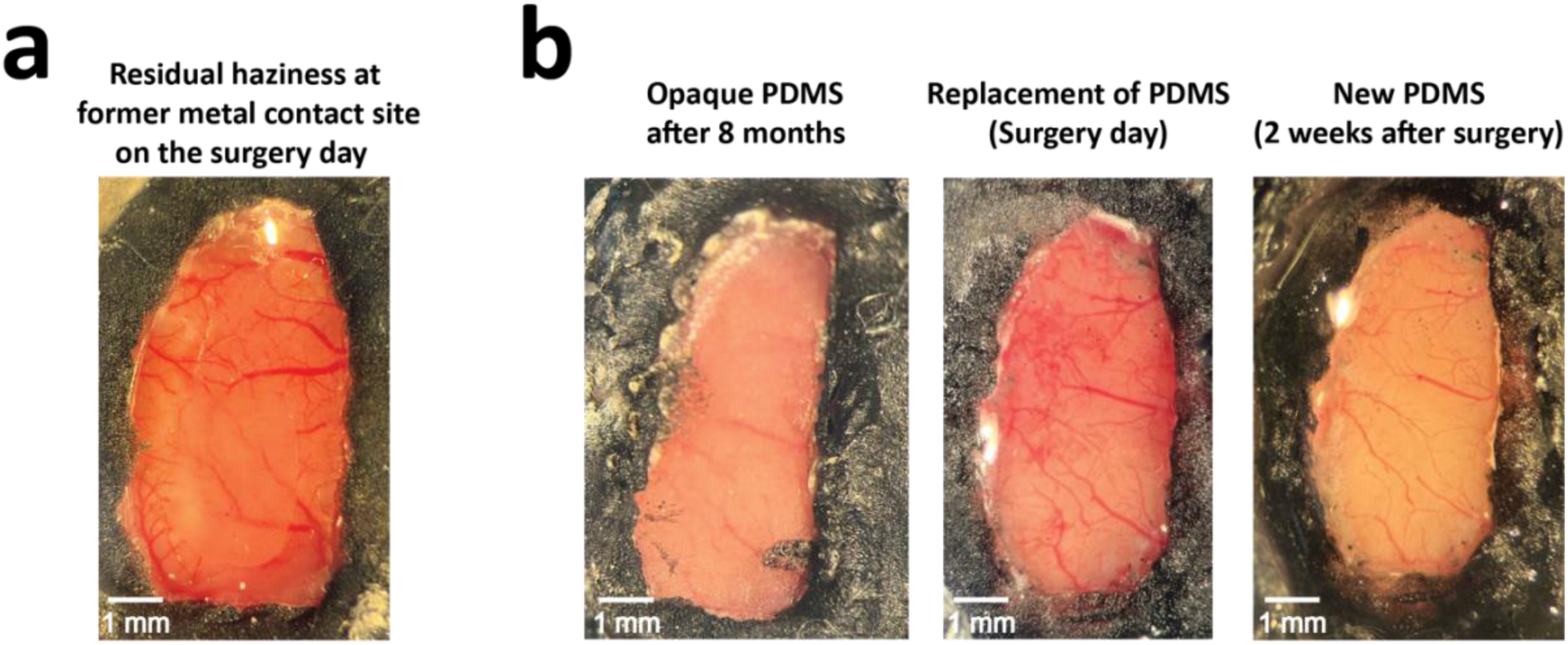
Potential complications following PDMS window placement. **(a)** An example of residual haziness in the posterior region of the PDMS window at the location of the former metal contact, observed on the surgery day (see Figure 2b9 and 10). **(b)** Replacement of an opaque PDMS window after 8 months. The left image shows an opaque PDMS window, the middle image shows the window immediately after replacement on the day of surgery, and the right image shows the clear brain following natural dissipation of residual blood after two weeks.

### Problem 3

PDMS loses its transparency after an extended period, usually several months (related to Step 2)

### Potential solution

PDMS typically maintains its clarity for about a year, although it may sometimes lose its transparency over time. If the window becomes excessively opaque (Fig. 7b, left), it can be replaced relatively easily through an additional surgical intervention. The procedure begins by drilling away the edge of the dental cement around the PDMS window. Once this is done, the PDMS can be removed using tweezers, making sure to remove any remaining dental cement waste. Incomplete removal of cement may cause the PDMS to tear during extraction, making the process more difficult. After the PDMS has been removed, the cranial window should be thoroughly cleaned to remove any remaining cement debris. Subsequently, one can follow the surgical steps detailed in Figure 2b.7-12 to insert a new window. Finally, the area is allowed to heal for 1-2 weeks, during which time any residual blood will naturally dissipate as shown in Fig. 7b, right.

### Problem 4

The animal’s breathing rate slows down when applying pressure to the PDMS with a metal plate (related to Step 2)

### Potential solution

If the animal experiences difficulty breathing during the surgical procedure when pushing the PDMS window, which could potentially be life-threatening, it indicates that too much force is being applied. In such cases, it is crucial to reduce the pressure and proceed with more care to avoid causing harm to the animal.

### Problem 5

An opening cannot be made in the PDMS window with precision, resulting in an uneven or poorly defined shape (Step 4)

### Potential solution

Creating an opening in the PDMS window can be difficult when using just a scalpel or needle, because of the material’s soft and pliable nature. This makes it challenging to achieve a precise incision. As of now, the most reliable method for performing this procedure is by using a biopsy needle, which offers greater control and precision when cutting through the PDMS window. It is recommended to avoid using extremely sharp instruments such as a scalpel or needle.

## Resource availability Lead contact

Further information and requests for resources and reagents should be directed to and will be fulfilled by the lead contact, Umberto Olcese.

## Technical contact

Technical questions on executing this protocol should be directed to and will be answered by the technical contact, Kengo Takahashi.

## Materials availability

All information on the materials utilized in this study is provided within this article. For further information, please reach out to the lead or technical contact.

## Data and code availability

Data presented in the paper will be made available on Zenodo upon publication.

## Acknowledgments

This project was made possible through the support of a grant from Templeton World Charity Foundation, Inc (TWCF0646) to RY, UO and CMAP, and of a grant from the European Union (EIC Pathfinder project NAP, grant agreement 101099310) to UO. The opinions expressed in this publication are those of the authors and do not necessarily reflect the views of Templeton World Charity Foundation, Inc. (https://www.templetonworldcharity.org/). The funders had no role in study design, data collection and analysis, decision to publish, or preparation of the manuscript. S.P. and R.Y. also acknowledge support from the NINDS (RM1NS132981) and NEI (R01EY035248).

We would also like to acknowledge administrative support by Reinder Dorman, technical assistance for PDMS production by Dr. Paola Fanzio, and the technical contribution of the Technology Center of the University of Amsterdam.

## Author contributions

Conceptualization, KT and UO.; methodology, KT and GJH.; investigation, KT, GJH, DB, JYW, and SPQ.; writing – original draft, KT.; writing – review and editing, KT, GJH, DB, JYW, SPQ, RY, CMAP, and UO.; funding acquisition, RY, CMAP, and UO.; supervision, UO.

